# PEMT: A patent enrichment tool for drug discovery

**DOI:** 10.1101/2022.07.28.501730

**Authors:** Yojana Gadiya, Andrea Zaliani, Philip Gribbon, Martin Hofmann-Apitius

## Abstract

**Motivation:** Drug discovery practitioners in industry and academia use semantic tools to extract information from online scientific literature to generate new insights into targets, therapeutics and diseases. However, due to complexities in access and analysis, patent-based literature is often overlooked as a source of information. As drug discovery is a highly competitive field, naturally, tools that tap into patent literature can provide any actor in the field an advantage in terms of better informed decision making. Hence, we aim to facilitate access to patent literature through the creation of an automatic tool for extracting information from patents described in existing public resources.

**Results:** Here, we present PEMT, a novel patent enrichment tool, that takes advantage of public databases like ChEMBL and SureChEMBL to extract relevant patent information linked to chemical structures and/or gene names described through FAIR principles and metadata annotations. PEMT aims at supporting drug discovery and research by establishing a patent landscape around genes of interest. The pharmaceutical focus of the tool is mainly due to the subselection of International Patent Classification (IPC) codes, but in principle, it can be used for other patent fields, provided that a link between a concept and chemical structure is investigated. Finally, we demonstrate a use-case in rare diseases by generating a gene-patent list based on the epidemiological prevalence of these diseases and exploring their underlying patent landscapes.

**Availability and implementation:** PEMT is an open-source Python tool and its source code and PyPi package are available at https://github.com/Fraunhofer-ITMP/PEMT and https://pvpi.org/project/PEMT/ respectively.

**Supplementary information:** Supplementary data are available at *Bioinformatics* online.

## 1. Introduction

Patents are an untapped source of scientific information which nevertheless play a vital role in reflecting the progress of organisations involved in drug discovery. Scientific content contained within patent documents can be overlooked due to the lengthy process involved in making the patent public, as compared to scientific publications which can be readily accessible to the community via pre-print servers. Moreover, the legal jargon used to describe the claims of the patent applications also introduces additional complexities in data analysis as compared to classical scientific publications. However, analysis of patents can be key in assessing and reviewing, for instance, a companies’ disease strategy (Roskams-Edris *et al*., 2019) or making well-informed target selection and prioritisation decisions (Jin and Wong, 2014).

Patent literature provides a different view of drug discovery by focusing on industry-specific genes or targets rather than scientific literature (Mucke, 2021). Databases such as PATENTSCOPE (https://www.wipo.int/patentscope/en/) and Espacenet (https://worldwide.espacenet.com/) specifically provide search and retrieval capabilities that cater to the specialized structured format of patent applications (Donald *et al*., 2018). Despite the intrinsic value in analysing all relevant patent data in a project, there is often a limit on the number of patents individual users can retrieve within any period of time. In addition, many existing resources have associated access fees, which may not be affordable for academia and small-scale companies. This situation gives rise to a need for open-source resources for analysing patent content.

Here, we present Patent EnrichMent Tool (PEMT), an automated patent extraction tool that takes advantage of information from patent databases and connects genes or chemicals to this information retrospectively. While patent officers are specifically trained in legal aspects of patent analysis, drug researchers have few open-source tools that enable them to perform a qualitative landscape evaluation on genes linked to certain diseases. PEMT is designed to provide them with such a tool. Furthermore, we demonstrate the applicability of this tool in the rare disease domain and show how the tool can assist the scientific community in exploring this untapped resource.

## 2. Methods and Material

### 2.1. Implementation details

PEMT is written in version-controlled software with Python 3.8 and can be found at PyPi (https://pypi.org/project/PEMT/) and on GitHub (https://github.com/Fraunhofer-ITMP/PEMT).

### 2.2. Data harmonization

PEMT deals with three types of entities: patents, genes, and modulators, such as chemicals or biological agents. Each of these entities are harmonized by mapping them to one or more well-known identifiers using cross-references available from existing biological databases. For genes, we leveraged the HUGO Gene Nomenclature Committee (HGNC) (Povey *et al*., 2001) database that enabled easy conversion of gene symbols or names to UniProt identifiers (UniProt Consortium, 2015). Similarly, to map gene identifiers to ChEMBL identifiers, and ChEMBL modulators to SureChEMBL identifiers (Papadatos *et al*., 2016), we made use of ChEMBL’s cross-reference system (Gaulton *et al*., 2012). Overall, genes were represented with UniProt and ChEMBL identifiers, modulators with ChEMBL and SureChEMBL identifiers, and patents with application numbers. The reasoning behind the selection of the abovementioned resources is discussed in **Supplementary Text 1.** This harmonization step served two main purposes: first, it increased the alignment of the underlying data resources with FAIR principles, and second, it allowed for data extraction from multiple resources in an efficient manner.

### 2.3. Design Architecture

PEMT takes a two-step approach to collect relevant patent documents (**Figure 1**). In the first step, chemical and biological modulators that directly regulate (i.e. activation or inhibition) specified genes of interest are extracted. For each gene of interest, a gene harmonization step described in the previous section is performed. Once we have the corresponding ChEMBL IDs for the gene, we query ChEMBL to identify experimentally validated compounds for the gene. Since ChEMBL has multiple approaches for flagging experimental data, we restricted the identified compounds to have binding or functional activity on the gene (**Supplementary Text 2**). Thus, the chemical extractor stage generated a pre-filtered and in-vivo validated list of chemical and biological modulators for the gene.

**Figure 1.**
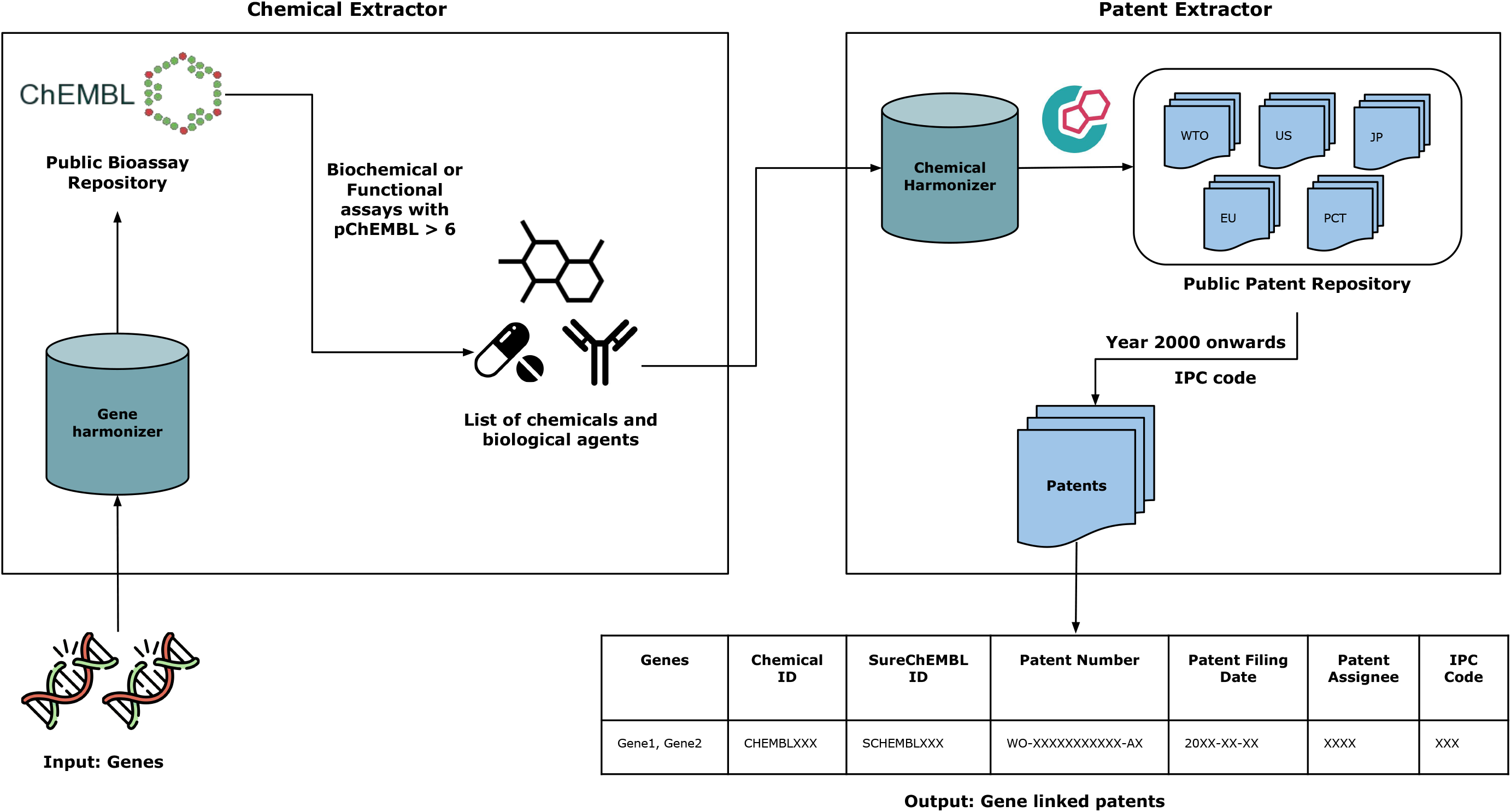
Overview of the framework of the Patent EnrichMent Tool (PEMT).

In the second stage, interlinking of identified modulators to patent documents is done by systematically querying SureChEMBL, a patent database. In order to extract relevant patents, first the chemical and biological modulators from the previous stage are harmonized to SureChEMBL IDs with the chemical harmonization. This is followed by querying the SureChEMBL database to extract all corresponding patent documents that reference the modulator. Simultaneously, a patent class based filtering was performed to retrieve patent documents playing a role in essential chemical or biological roles. (**Supplementary Text 3**). PEMT recognizes each patent document as unique based on its patent number which consists of a country code (e.g. WO/US/EP), a 7-11 digit number, and the patent grant status number (e.g. A1, B2, etc) each separated by a dash (-). Together, these two steps facilitate the linking of patent documents to genes of interest via chemicals or biological modulators.

## 3. Case scenario

### 3.1. Data generation

Orphanet (www.orpha.net) is a service catalog of rare disease data which includes the Orphanet Ontology of Rare Diseases (ORDO), as well as information on the epidemiological occurrence and the biological mechanisms (disease-gene interactions) involved in rare diseases (Weinreich *et al*., 2008). We made use of two data categories from Orphanet, i.e. epidemiological data and biological mechanisms, to demonstrate the applicability of PEMT.

As a starting point, we selected a subset of genes based on their corresponding disease prevalence from the current list of 3,886 diseases and 2,637 genes available in Orphanet. We also filtered out the diseases with an incidence rate lower than 10. This selection criterion resulted in 59 diseases and 56 genes, representing the most common of the rare diseases, where significant patenting activity would be reasonably anticipated to have occurred in the past. These 56 genes were then provided as inputs for the PEMT tool.

### 3.2. Data analysis

PEMT generated connections between genes to patents via modulators. Based on publicly available bioassays, only 12 genes were associated with modulators (**Supplementary Figure 1**). The number of agents ranged between 115 (for FGFR3) and 1 (for MECP2) and a total of 469 were extracted from this step. A further reduction in the chemical space (of the 469 chemicals) was performed, based on their patentability, resulting in 143 modulators. From these 143 modulators, only 97 were identified to be linked to pharmaceutically relevant patents (**Supplementary Table 2**). Ultimately, 135 unique patents belonging to 10 genes (2 did not have any patent information connected to them), linked via 97 modulators, were extracted using the PEMT tool (**Supplementary Figure 2**).

Patents covered two geographical jurisdictions: 83 from the United States of America and 19 from the European Patent Office (EPO). Furthermore, 106 patents are not yet granted while 29 patents have already been granted (**Supplementary Table 3**). Of the 106 non-granted patents, we found that 33 patents were from the World Intellectual Property Organization (WIPO) which includes patent applications but does not grant them, unlike EPO or USPTO who have patent granting authority. We also found that these patents cover a range of different cooperative patent classification (CPC) codes: C07D (82 patents), C07K (18 patents), C12Q (14 patents), C07H (4 patents), A61P (9 patents), C12P (3 patents), C07F (5 patents), and C07C (1 patent) and are distributed over the year 2000-2021 (**Supplementary Figure 3**). Moreover, upon assessing the patent assignees, we found 85 patents belonged to pharmaceutical industries, 23 patents to academic institutes, and 25 patents to individuals.

Lastly, we looked at the historical timeline for each patented gene to understand the genes’ importance over the patent landscape (**Supplementary Figure 4**). FGFR3, one of the genes that received major attention between 2002 and 2008, was found to be associated with multiple diseases. In the beginning of this time period, FGFR3 was found to be associated with Muenke syndrome which had no interventional clinical studies, leading to a peak in patenting activity. Despite these efforts, only a single completed clinical trial in 2005 (NCT00106977) was performed that aimed at better understanding disease aetiology. Later, in 2004, it was found that FGFR3 played a role in carcinomas, such as multiple myeloma and bladder carcinoma (Grand *et al*., 2004) and interest within the industry to understand the effect of this target in carcinoma remained. Nonetheless, only a marginal efficacy was demonstrated in this case as well (Chae *et al*., 2017) and in the following years, interest in the target began to wane. Conversely, other targets, including SLC2A3 began surfacing in patents in 2015 and contributed to the highest number of patents, primarily due to their association with Huntington’s disease, a prominent neurodegenerative disease.

## 4. Discussion and Future Work

The Patent EnrichMent Tool (PEMT) is an open-source and easy-to-use tool providing the scientific community with the opportunity to extract pharmaceutically relevant patent documents for genes of interest. This efficient linking of genes to patents could potentially open a link to enrich existing biomedical knowledge graphs with unmined patent literature. Moreover, PEMT can help in providing an overview of the target prioritization shift over time. Hence, with the PEMT we hope to enhance the accessibility and applicability of patent literature within the scientific community.

## Acknowledgement and Funding

This research was funded by BMBF TreatKCNQ, grant no. 01GM2003A. The authors thank Daniel Domingo-Fernández, Sarah Mubeen, and the anonymous reviewers for their comments and suggestions on the manuscript.

---

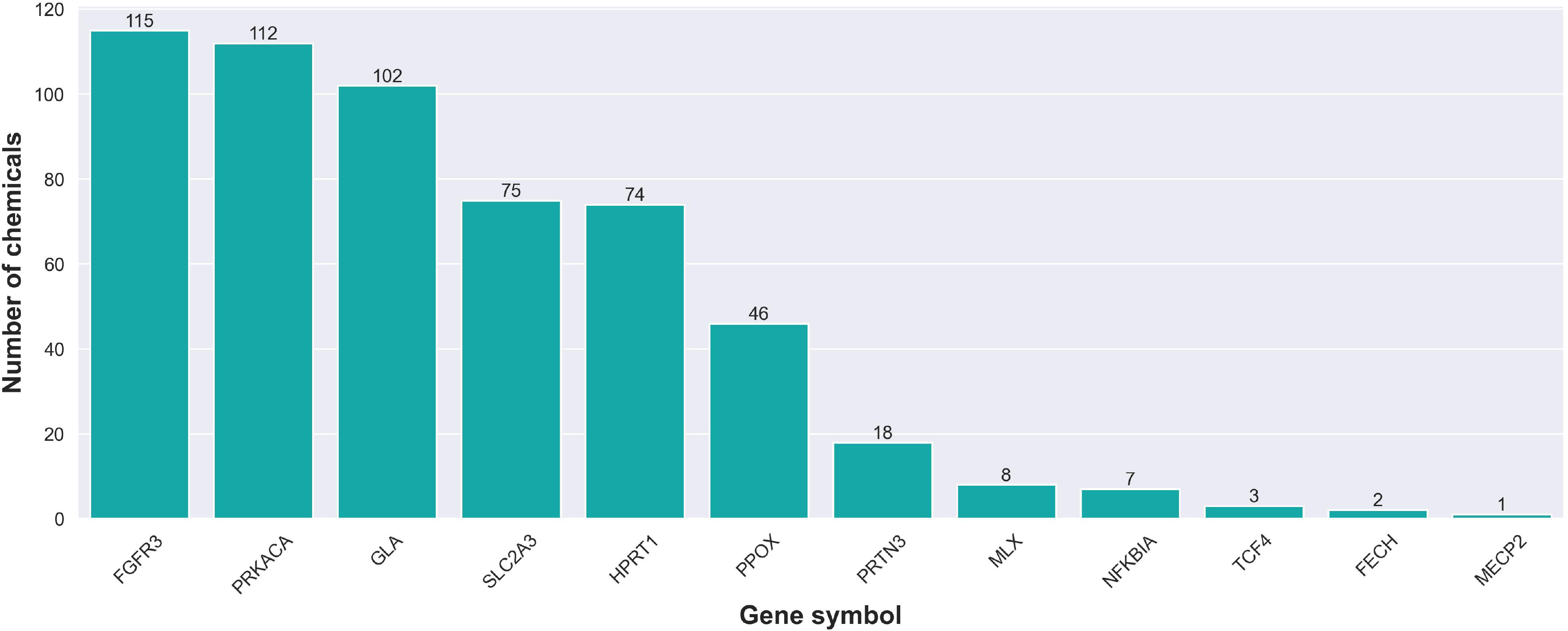

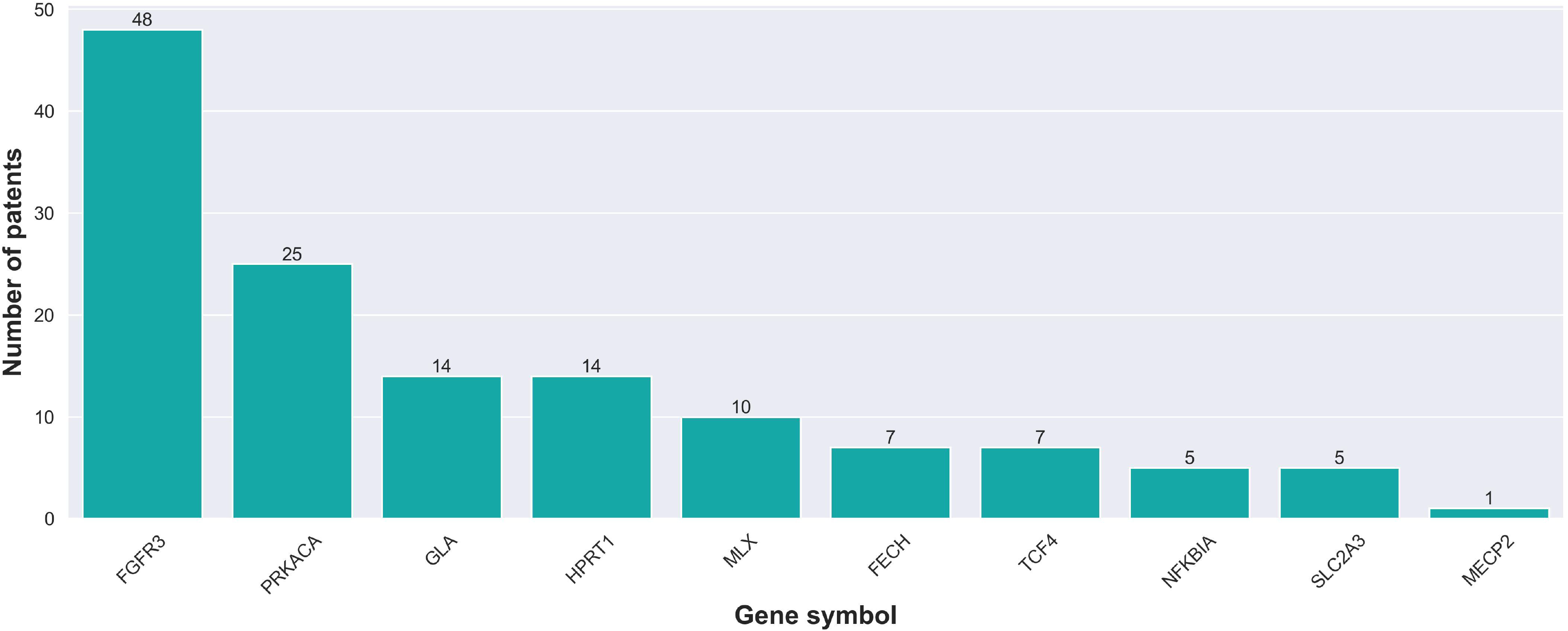

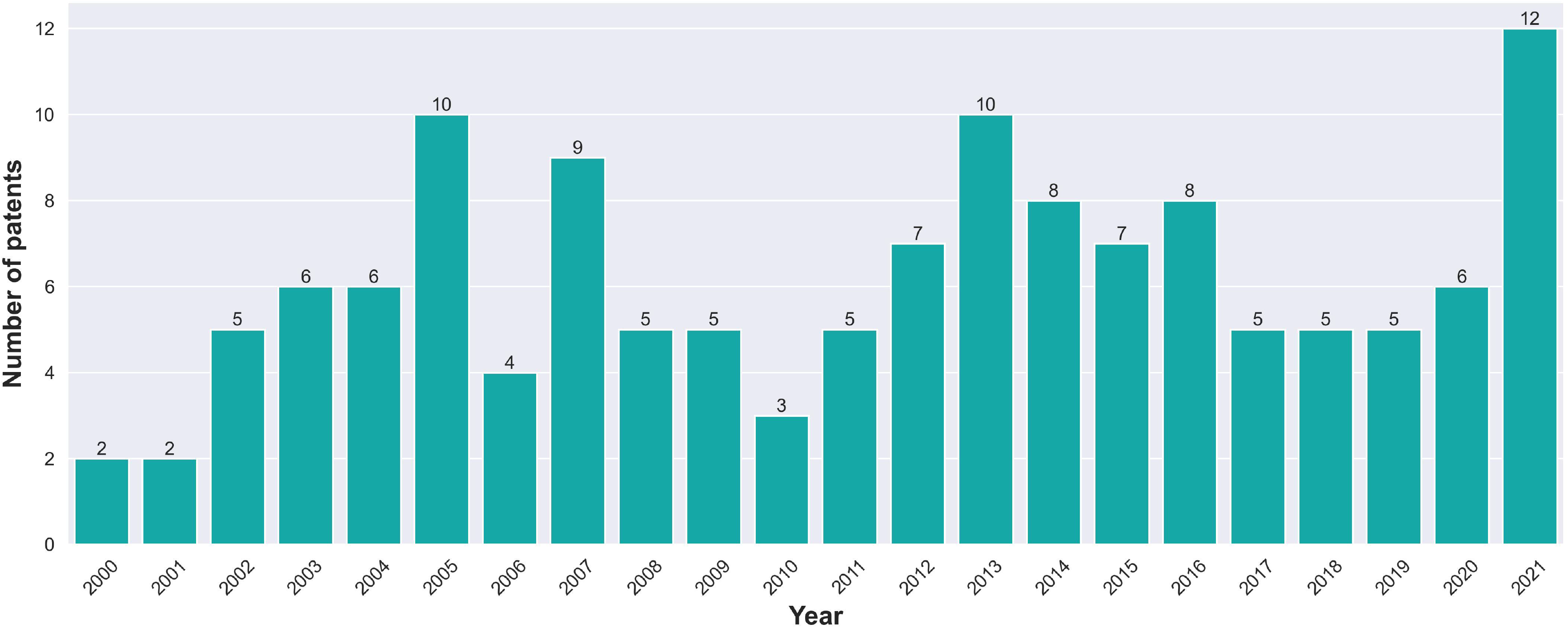

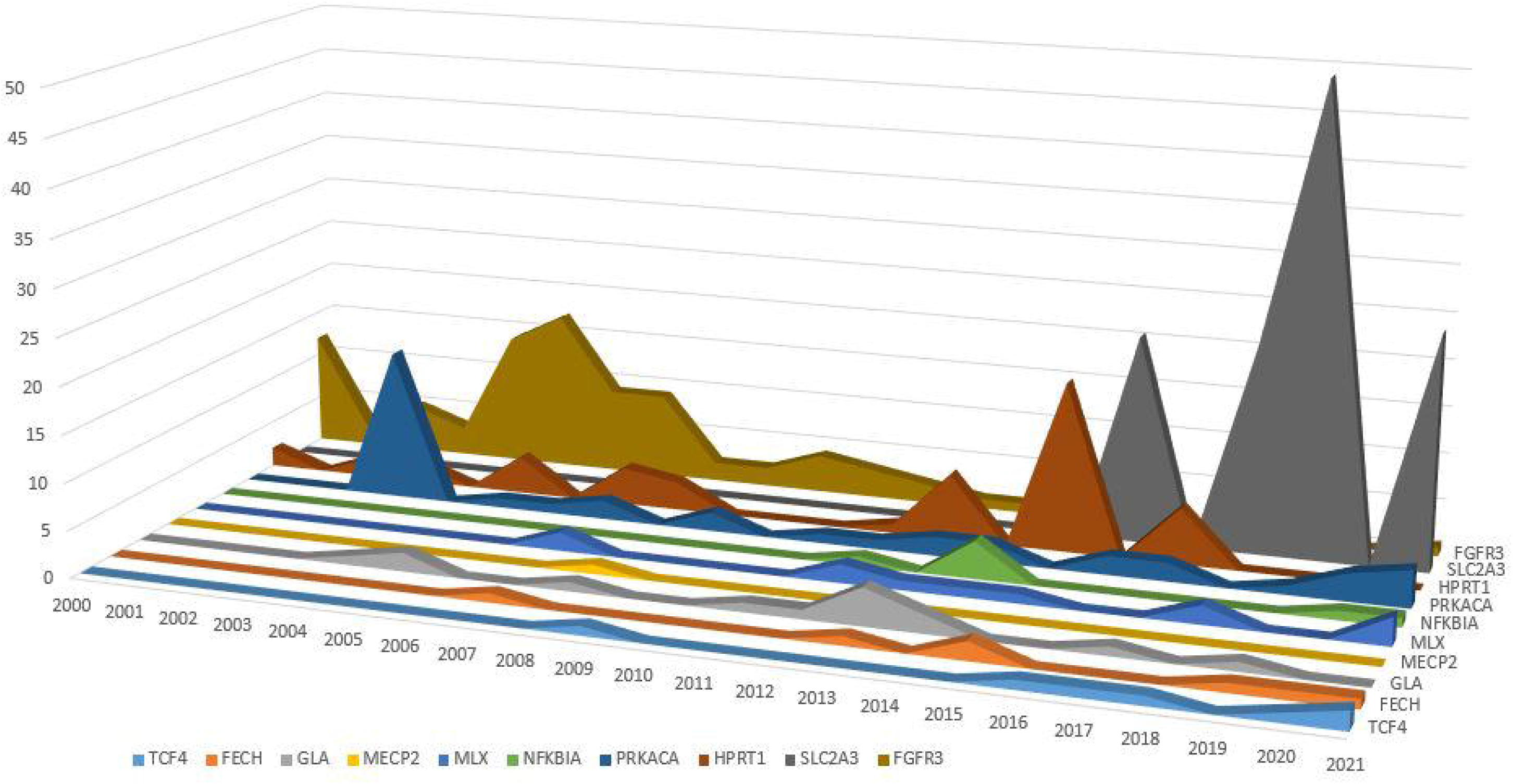

