## Supplementary file for "PEMT: A patent enrichment tool for drug discovery"

### Supplementary Material to “PEMT: A patent enrichment tool for drug discovery”

**SUPPLEMENTARY TEXT**

**Supplementary Text 1: Reason for selection of chemical and patent data resources**

ChEMBL: ChEMBL (Gaulton *et al.*, 2012) is one of the major public resources for chemical biology. ChEMBL version 30 contains 2.2 million compounds and 1.5 million assays belonging to biochemical, cellular, and/or phenotypic nature. Moreover, human-in-the-loop curation of the ingested data ensured the quality of the data. Given the vast amounts of data points that enabled the linking of chemical entities to assay data, ChEMBL was used as the source for extraction of bioassay data.

SureChEMBL: SureCheMBL (Papadatos *et al.*, 2016) is a public patent data resource that explicitly collects patents from several patent authorities such as the World Intellectual Property Organisation (WIPO) and the United States Patent and Trademark Office (USPTO) among others. Furthermore, SureChEMBL makes use of a proprietary text mining and image recognition system that enables the data within patents to be parsed allowing for interlinking biological entities, such as chemicals and patents. As SureChEMBL captures patents globally, it was the patent extraction source used for PEMT.

UniProt and HGNC: UniProt (UniProt Consortium, 2015) and HUGO Gene Nomenclature Committee (HGNC) (Povey *et al.*, 2001) are two of the largest protein data resources that currently exist. Moreover, given their extensive cross-referencing features with chemical databases such as ChEMBL, they were a suitable choice for the representation of genes or proteins within PEMT.

**Supplementary Text 2: Filtering approach for selecting chemical-gene pairs**

The bioassay data in ChEMBL were classified based on their activity as functional, biochemical, absorption, distribution, metabolism, and excretion (ADME), toxicity, and physicochemical. Since PEMT is designed to extract bioactive chemicals that have a causal effect on the gene of interest, it restricted the experimental search space to biochemical assays that were either functional or binding. Thus, chemical-gene pairs with a high confidence score were selected. A confidence score is a number between 1 to 9 which is manually assessed by the ChEMBL team based on the assay or experiment type (lower scores are assigned to cellular/phenotypic assay types and higher scores are assigned to biochemical ones). Furthermore, each bioassay experiment was assigned a pChEMBL value, which is mathematically expressed as the negative logarithmic of the molar concentration, and directly correlated with the activity of the given chemical. Thus, PEMT also leveraged this metric and filtered chemicals that had a pChEMBL > 6 (i.e. those chemicals that have activity in the submicromolar range).

**Supplementary Text 3: Filtering approach for selecting patents**

The patent list retrieved from SureChEMBL was filtered based on the following two aspects: patent validity (i.e., patents that are still active and have not yet expired) and pharmacological relevance (i.e. those patent documents that correspond to or underline any relevant physiological change to the human body). This filtering ensured that the resultant patent was both pharmaceutically relevant and valid. It is well-known that a patent is valid for 25 years; hence, PEMT made use of this timeline to identify patents that are still active by collecting patents registered from 2000 onwards. The pharmaceutically relevant patents were identified based on their International Patent Classification (IPC) code (<https://www.wipo.int/classifications/ipc/en/>). The IPC code is a classification tree representing the content coverage of a patent and hence each patent can belong to one or more IPC codes. We manually filtered 34 IPC codes based on their relevance in drug discovery **(Supplementary Table 1)**. Within these 34 IPC codes, 6 classes: A23, A61, C01, C07, C08, and C12 were covered. These classes include information on usage for chemicals for medicinal purposes and chemical formulation information, thus covering the chemobiology domain. We omitted A61K code which covered synthetic preparation for medical, dental, and hygiene purposes as patent documents with this IPC code usually involve synthesis of chemicals that may not be specific for health or pharmacological use and hence may not specify any details on gene target effects within humans. This omission restricts PEMT results to pre-clinical research stages or to repurposing of known drugs to novel targets.

**SUPPLEMENTARY FIGURES**

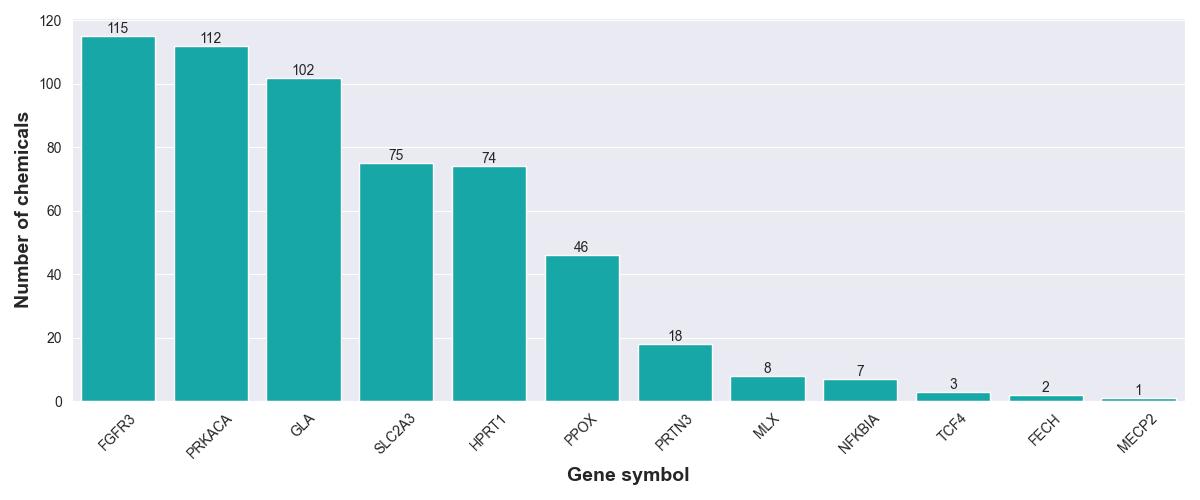
**Supplementary Figure 1**: Plot indicating the number of chemicals and biological agents extracted from bioassays for genes of interest (HGNC gene symbols). The genes of interest in this case are the rare disease-targeting gene from Orphanet.

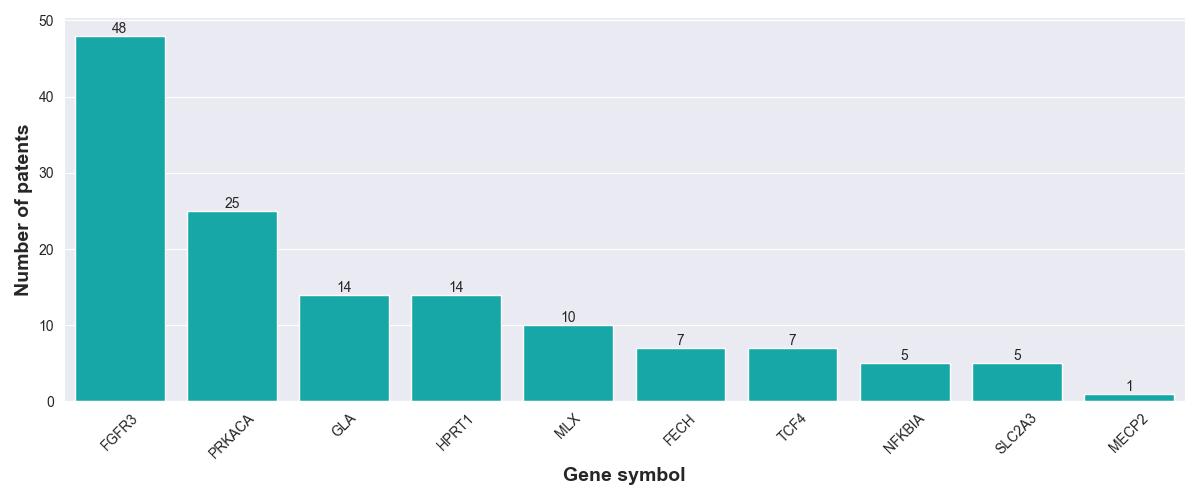
**Supplementary Figure 2**: Plot indicating the number of patents linked with genes of interest (HGNC gene symbols) based on PEMT.

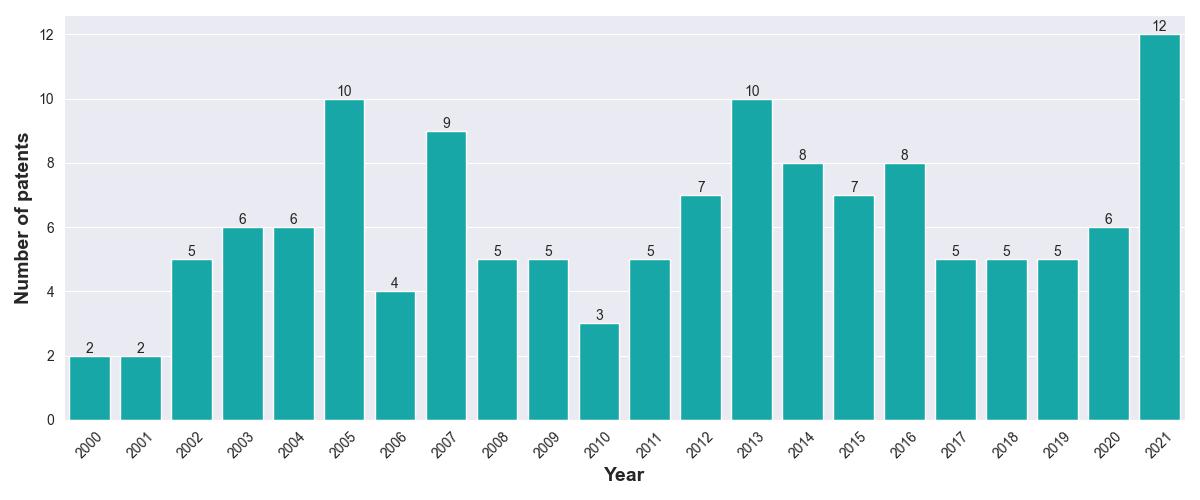
**Supplementary Figure 3**: Distribution of the extracted patent landscape from 2000 - 2021.

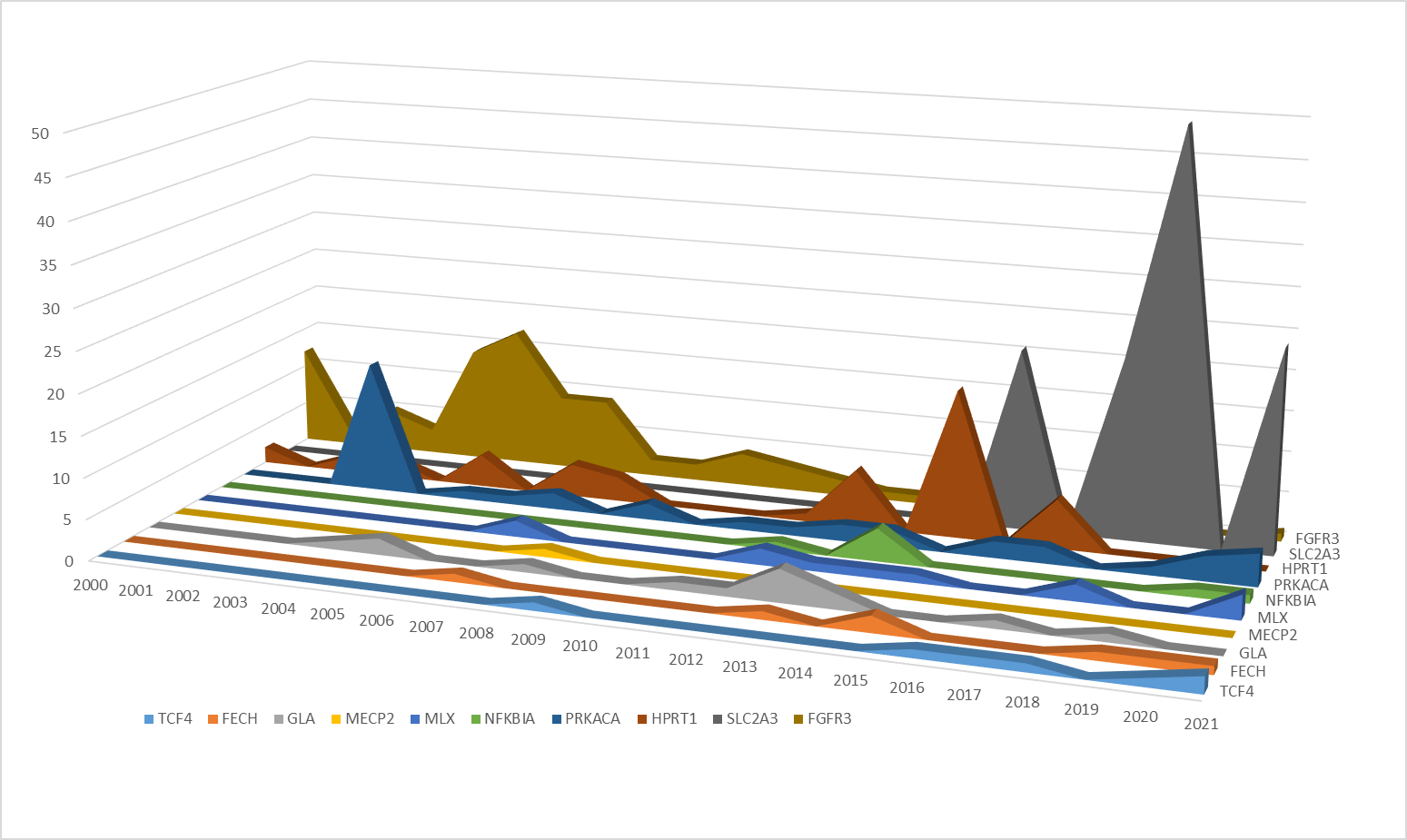

**Supplementary Figure 4**: Historical distribution of the patents for each gene from 2000 - 2021.

**SUPPLEMENTARY TABLES**

| **IPC Code** | **Description of the class** |
| --- | --- |
| A23B | Modulators relevant for preserving purposes |
| A23C | Modulators containing protein compositions from dairy products |
| A23D | Modulators correlating to edible oils and fats |
| A23F | Modulators derived from coffee, tea, and their substitutes |
| A23G | Cocoa related modulators |
| A23J | Modulators with effect on protein compositions in food |
| A23L | Modulators found in food, foodstuff, or non-alcoholic beverages |
| A23N | Modulators involved in treating fruits, vegetables, flowers |
| A23P | Modulators found in food |
| A61B | Modulators involved in medical, veterinary, or in hygiene science |
| A61C | Modulators used for diagnosis or in surgery |
| A61D | Modulators used in veterinary instruments and tools |
| A61P | Modulators used in medicinal preparations for specific therapeutic activities |
| C01B | Modulators containing non-metallic elements |
| C01C | Modulators containing ammonia and/or cyanogen |
| C01G | Modulators containing metallic elements |
| C07B | Modulators belonging to organic chemistry |
| C07C | Acyclic or carbocyclic modulators |
| C07D | Heterocyclic modulators |
| C07F | Acyclic, carbocyclic or heterocyclic modulators with elements other than carbon, hydrogen, halogen, oxygen, nitrogen, sulphur, selenium, or tellurium |
| C07G | Modulators with unknown constitutions |
| C07H | Sugar derived modulators |
| C07J | Modulators belonging to steroids |
| C07K | Modulators containing peptides |
| C08B | Modulators containing polysaccharides and its derivatives |
| C08C | Modulators involved in modification of rubber |
| C08F | Macromolecular modulators obtained by reactions between carbon-carbon unsaturated bonds |
| C08G | Macromolecular modulators obtained by other reactions than C08F |
| C08H | Natural macromolecular derivative compounds |
| C08J | Modulators that do not belong to classes C08B, C08C, C08F, C09G, or C08H |
| C08K | Modulators containing inorganic or non-macromolecular organic substrates |
| C08L | Compositions of macromolecular modulators |
| C12P | Modulators requiring fermentation or enzyme for synthesis |
| C12Q | Modulators involved in testing of enzymes, nucleic acids, or microorganisms |

**Supplementary Table 1**: The 34 IPC codes along with their description that are used for filtering patents in PEMT.

| **ChEMBL identifiers of modulators** | **Number of chemicals** | **Number of patents** |
| --- | --- | --- |
| CHEMBL379218 | 1 | 19 |
| CHEMBL49120 | 1 | 15 |
| CHEMBL2063869 | 1 | 10 |
| CHEMBL1233663, CHEMBL299763 | 2 | 9 |
| CHEMBL195008, CHEMBL1233603 | 2 | 8 |
| CHEMBL254381, CHEMBL1944698 | 2 | 7 |
| CHEMBL45827 | 1 | 6 |
| CHEMBL193306, CHEMBL4632920, CHEMBL4645739, CHEMBL4640792, CHEMBL4649871, CHEMBL4645691, CHEMBL4641999, CHEMBL195903, CHEMBL195398, CHEMBL195847, CHEMBL194498, CHEMBL4633651, CHEMBL4635844, CHEMBL4649877, CHEMBL4634839, CHEMBL4634395, CHEMBL4649211, CHEMBL4637272, CHEMBL4637311, CHEMBL4647586, CHEMBL4638502, CHEMBL4638556, CHEMBL4635864, CHEMBL4637262, CHEMBL4635564, CHEMBL4637134, CHEMBL4647311 | 27 | 5 |
| CHEMBL1952210, CHEMBL3908855, CHEMBL1952211, CHEMBL606964 | 4 | 4 |
| CHEMBL4648466, CHEMBL4648473, CHEMBL2153485, CHEMBL2153483, CHEMBL2153497, CHEMBL140808, CHEMBL143147, CHEMBL301612, CHEMBL2153480, CHEMBL2153478, CHEMBL2153477, CHEMBL2153476, CHEMBL4633530 | 13 | 3 |
| CHEMBL374805, CHEMBL228133, CHEMBL3613599, CHEMBL226625, CHEMBL438483, CHEMBL2153481, CHEMBL2153484, CHEMBL272552, CHEMBL271669, CHEMBL429586, CHEMBL213618, CHEMBL2063868, CHEMBL378963, CHEMBL299194, CHEMBL383264, CHEMBL383541, CHEMBL207544, CHEMBL1507269, CHEMBL469844 | 19 | 2 |
| CHEMBL4282830, CHEMBL4280490, CHEMBL4291117, CHEMBL4277753, CHEMBL75188, CHEMBL299026, CHEMBL4290716, CHEMBL57347, CHEMBL57323, CHEMBL520851, CHEMBL4285691, CHEMBL432738, CHEMBL2414636, CHEMBL56236, CHEMBL57366, CHEMBL301483, CHEMBL51028, CHEMBL51485, CHEMBL298679, CHEMBL50647, CHEMBL1482983, CHEMBL1429822, CHEMBL227605, CHEMBL4071045 | 24 | 1 |

**Supplementary Table 2**: Overview of the modulators linked to patents along with their respective patent counts.

| **Patent status** | **Patent designation code** | **Number of patents** |
| --- | --- | --- |
| Granted patents | A1 document | 90 |
|  | A2 document | 13 |
|  | A4 document | 2 |
| Patents in the process of being granted | B1 document | 3 |
|  | B2 document | 26 |

**Supplementary Table 3**: Overview of the current status of patents within the market. The patent status indicates whether the patents are still in the process of being granted or have already been granted. Since these designations are specific with respect to the European Patent Office (EPO), the definitions for each of these can be found on the EPO website (<https://www.epo.org/searching-for-patents/helpful-resources/first-time-here/definitions.html>).
